# Deep neural networks: a new framework for modelling biological vision and brain information processing

**DOI:** 10.1101/029876

**Authors:** Nikolaus Kriegeskorte

**Affiliations:** Medical Research Council Cognition and Brain Sciences Unit, Cambridge, UK

**Keywords:** biological vision, computer vision, object recognition, neural network, deep learning, artificial intelligence, computational neuroscience

## Abstract

Recent advances in neural network modelling have enabled major strides in computer vision and other artificial intelligence applications. Human-level visual recognition abilities are coming within reach of artificial systems. Artificial neural networks are inspired by the brain and their computations could be implemented in biological neurons. Convolutional feedforward networks, which now dominate computer vision, take further inspiration from the architecture of the primate visual hierarchy. However, the current models are designed with engineering goals and not to model brain computations. Nevertheless, initial studies comparing internal representations between these models and primate brains find surprisingly similar representational spaces. With human-level performance no longer out of reach, we are entering an exciting new era, in which we will be able to build neurobiologically faithful feedforward and recurrent computational models of how biological brains perform high-level feats of intelligence, including vision.

## INTRODUCTION

The brain is a deep and complex recurrent neural network. However, the computational models of brain information processing that have dominated computational neuroscience, in vision and beyond, are largely shallow architectures performing simple computations. Unsurprisingly, complex tasks such as visual object recognition have remained beyond the reach of computational neuroscience. In this paper, I argue that recent advances in neural network models (LeCun et al. 2015) will usher in a new era of computational neuroscience, in which we will engage real-world tasks that require rich knowledge and complex computations.

Neural networks are an old idea, so what is new now? Indeed, the history of neural networks is roughly coextensive with that of modern computing machines. John von Neumann and Alan Turing, whose ideas shaped modern computing technology, both explored network models inspired by the brain.

An early mathematical model of a single neuron was suggested by McCulloch & Pitts (1943). Their binary threshold unit computed a weighted sum of a number of inputs, and imposed a binary threshold, implementing a linear discriminant. Responding to a pattern of continuous inputs with a single binary output, the threshold unit provided an intuitive bridge between the biological hardware of a spiking neuron and categorisation, a hallmark of cognition.

Discriminating categories that are not linearly separable in the input requires an intervening layer of nonlinear transformations between the input and the output units. It took a while until the field found ways to automatically train such multi-layer networks with input-output pairs. The most influential solution to this problem is the backpropagation algorithm, a gradient-descent method that makes iterative small adjustments to the weights so as to reduce the errors of the outputs (Werbos 1981; Rumelhart et al. 1986).

Backpropagation led to a second wave of interest in neural networks in cognitive science and artificial intelligence (AI) in the 1980s. In cognitive science, neural network models of toy problems boosted the theoretical notion of parallel distributed processing (Rumelhart & McClelland 1988). However, backpropagation models did not work well on complex real-world problems such as vision. Less brain-inspired models, using hand-engineered representations and machine learning techniques including support vector machines, appeared to provide better engineering solutions for computer vision and AI. As a consequence, neural networks fell out of favour in the 1990s.

Despite a period of disenchantment among the wider brain and computer science communities, neural network research has an unbroken history (Schmidhuber 2014) in both theoretical neuroscience and computer science. It was pursued throughout the 1990s and 2000s by a smaller community of scientists who didn’t relent because they realised that the difficulties encountered were not fundamental limitations of the approach, but merely high hurdles to be overcome through a combination of better learning algorithms, better regularisation, and larger training sets – with computations boosted by better computer hardware. Their efforts have been fruitful.

How previous attempts to understand complex brain information processing fell short

The cognitive and brain sciences have gone through a sequence of transformations, with different fields dominating each period. Cognitive psychology attempted to illuminate behaviourism’s black box with theories of information processing. However, it lacked fully explicit computational models. Cognitive science made information processing theory fully explicit, but lacked constraints from neurophysiological data. This made it difficult to adjudicate between alternative models consistent with behavioural data. Connectionism within cognitive science offered a neurobiologically plausible computational framework. However, neural network technology was not sufficiently advanced to take on real-world tasks such as object recognition from photographs. As a result, neural networks did not initially live up to their promise as AI systems and in cognitive science, modelling was restricted to toy problems. Cognitive neuroscience brought neurophysiological data into investigations of complex brain information processing. However, our hands full with the challenges of analysing the new and unprecedentedly rich brain imaging data, our theoretical sophistication initially slipped, back to the stage of cognitive psychology, and we began (perhaps reasonably) by mapping box-and-arrow models onto brain regions. Computational neuroscience uses computational models to predict neurophysiological and behavioural data. However, at this level of rigour we have not been able to engage complex real-world computational challenges and higher-level brain representations. Now deep neural networks provide a framework for engaging complex cognitive tasks and predicting both brain and behavioural responses.

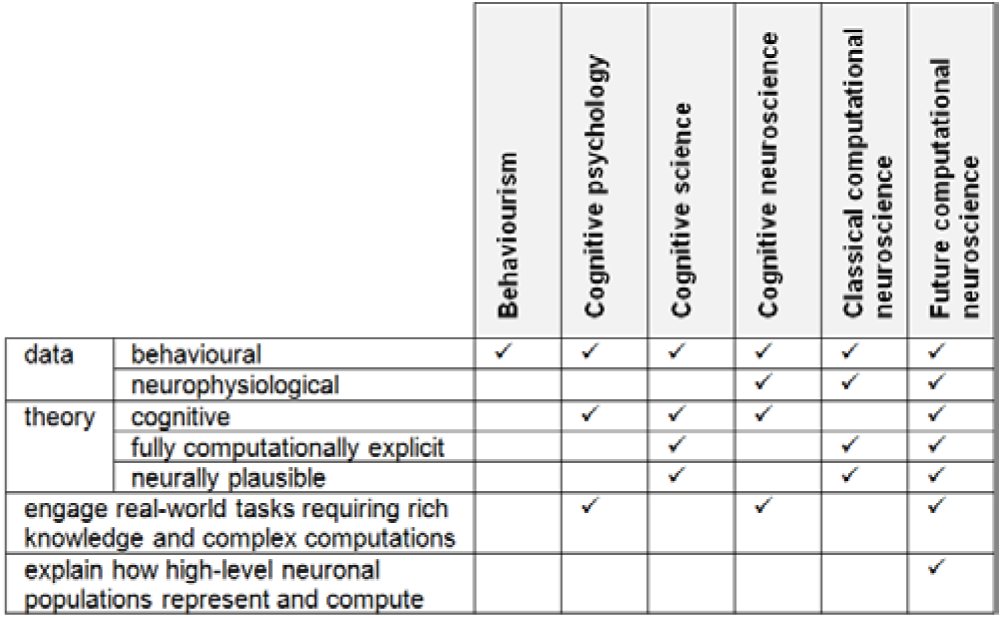

In the last few years, neural networks have finally come into their own. They are currently conquering several domains of AI, including the hard problem of computer vision.

Computer vision competitions like ImageNet (Deng et al. 2009) use secret test sets of images, providing rigorous evaluations of performance. In 2012, the ImageNet classification competition was won by a large margin by a neural net model built by Krizhesvsky et al. (2012), whose deep convolutional architecture enabled a leap in performance. Human performance levels, although still superior, suddenly did not seem entirely unattainable for computer vision any longer – at least in restricted domains like visual object classification. Krizhesvsky et al. (2012) marked the beginning of the dominance of neural networks in computer vision. In the last three years, error rates have dropped further roughly matching human performance in the restricted domain of visual object classification. Neural nets have also recently been very successful in other domains, including speech recognition (Sak et al. 2014) and machine translation (Sutskever et al. 2014).

AI has entered an era in which systems directly inspired by the brain dominate practical applications. The time has come to bring this brain-inspired technology back to the brain. We are now in a position to integrate neural network theory with empirical systems neuroscience and to build models that engage the complexities of real-world tasks, use neurobiologically plausible computational mechanisms, and predict neurophysiological and behavioural data.

The theoretical and engineering developments are progressing at an unprecedented pace. Many of the insights currently gained in engineering are likely to be relevant for brain theory. Recent methods for comparing internal representations in neural population codes between models and brains enable us to test models as theories of brain information processing (Dumoulin & Wandell 2008; Kay et al. 2008; Kriegeskorte et al. 2008a,b, 2013; Kriegeskorte 2011; Mitchell et al. 2008; Nili et al. 2014).

This paper serves to introduce a broad audience of vision and brain scientists to the computational literature on neural networks, including some of the recent advances of this modelling framework in engineering, and to review the first few studies using such models to explain brain data. What emerges is a new framework for bringing computational neuroscience to high-level cortical representations and complex real-world tasks.

The following *Primer* section introduces the basics of neural network models, including their learning algorithms and universal representational capacity. The section *Feedforward neural networks for visual object recognition* describes the specific large-scale object recognition networks that currently dominate computer vision and discusses what they share and do not share with biological vision systems. The section *Early studies testing deep neural nets as models of biological brain representations* reviews the first few studies that have begun to empirically compare internal representations between artificial neural nets and biological brains. The section *Recurrent neural nets for vision* describes networks using recurrent computation. Recurrence is an essential component of biological brains, might implement inference on generative models of the formation of the input image, and represents a major frontier for computational neuroscience. Finally, the *Conclusions* section considers critical arguments, upcoming challenges, and the way ahead toward empirically justified models of complex biological brain information processing.

## A PRIMER ON NEURAL NETWORKS

### A unit computes a weighted sum of its inputs and activates according to a nonlinear function

We will refer to model “neurons” as *units,* in order to maintain a distinction between the biological reality and the highly abstracted models. The perhaps simplest model unit is a linear unit, which outputs a linear combination of its inputs (Fig. 1a). Such units, combined to form networks, can never transcend linear combinations of the inputs. This is illustrated in Fig. 2b, which shows how an output unit linearly combining intermediate-layer linear-unit activations is just adding up ramp functions, and thus computing a ramp function. A multi-layer network of linear units is equivalent to single-layer network whose weights matrix W’ is the product of the weights matrices W_i_ of the multi-layer network. Nonlinear units are essential because their outputs provide building blocks (Fig. 2c) whose linear combination one level up enables us to approximate any desired mapping from inputs to outputs, as we will see in the next section.

**Figure 1:**
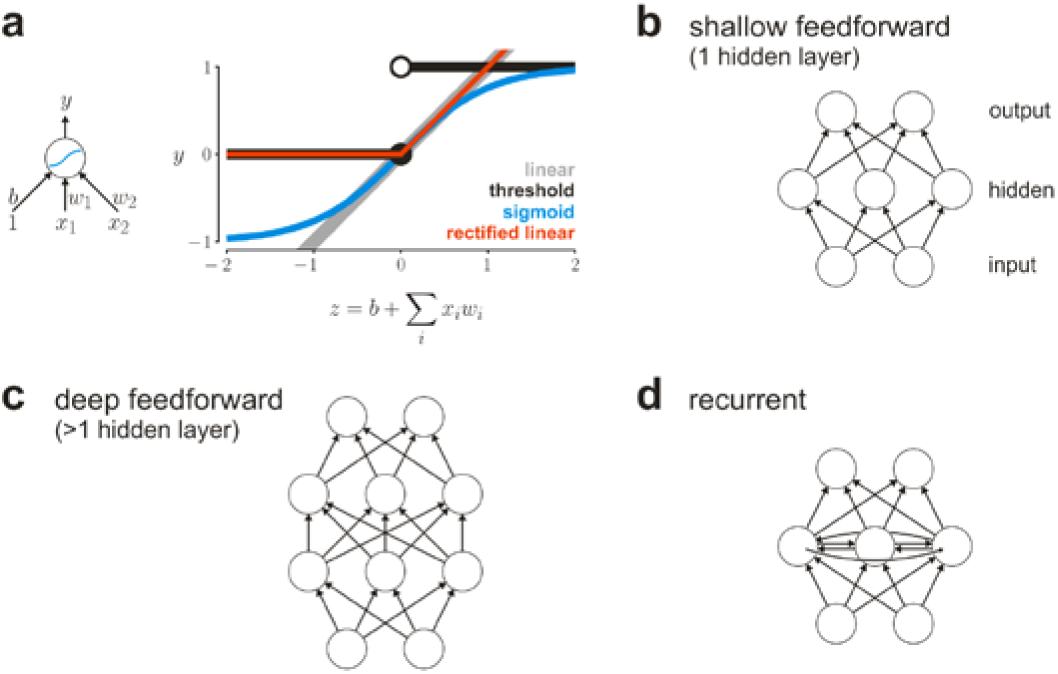
Artificial neural networks: basic units and architectures. A typical model unit (a, left) computes a linear combination z of its inputs x_i_ using weights w_i_ and adding a bias b. The output y of the unit is a function of z, known as the *activation function* (a, right). Popular activation functions include linear (gray), theshold (black), sigmoid (hyperbolic tangent shown here, blue), and rectified linear (red) functions. A network is referred to as *feedforward* (b, c) when its directed connections do not form cycles and as *recurrent* (d) when they do form cycles. A *shallow* feedforward net (b) has zero or one hidden layers. Nonlinear activation functions in hidden units enable a shallow feedfoward net to approximate any continuous function (with the precision depending on the number of hidden units). A *deep* feedforward net (c) has more than one hidden layer. Recurrent nets generate ongoing dynamics, lend themselves to the processing of temporal sequences of inputs, and can approximate any dynamical system (given a sufficient number of units).

**Figure 2:**
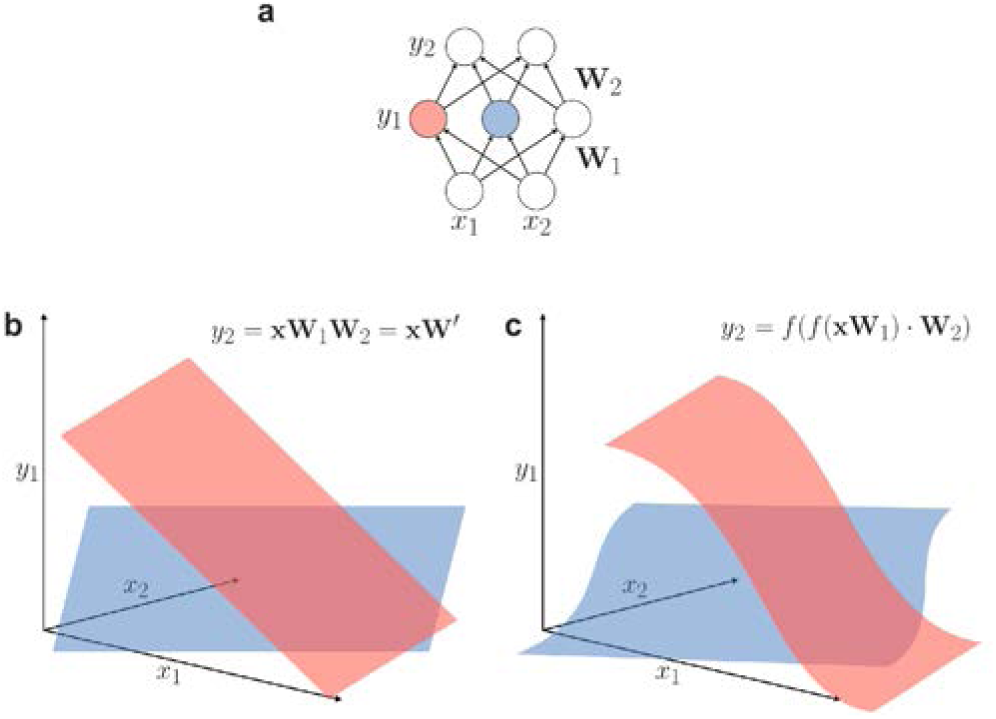
Networks with nonlinear hidden units can approximate arbitrary nonlinear functions. (a) Feedforward neural net with a single hidden layer. (b) Activation of the pink and blue hidden units as a function of the input pattern (x_1_, x_2_) when the hidden units have *linear* activation functions. Each output unit (y_2_) will compute a weighted combination of the ramp-shaped (i.e. linear) activations of the hidden units. The output, thus, remains a linear combination of the input pattern. A linear hidden layer is not useful because the resulting network is equivalent to a linear network without a hidden layer intervening between input and output. (c) Activation of the pink and blue hidden units when these have sigmoid activation functions. Arbitrary continuous functions can be approximated in the output units (y_2_) by weighted combinations of a sufficient number of nonlinear hidden-unit outputs (y_1_).

A unit in a neural net uses its input weights w to compute a weighted sum z of its input activities x and passes the result through a (typically monotonic) nonlinear function f to generate the unit’s activation y (Fig. 1a). In early models, the nonlinearity was simply a threshold (McCulloch & Pitts 1943; Rosenblatt 1958; Minsky & Papert 1972), making each unit a binary linear discriminant. For a single threshold unit, the perceptron learning algorithm provides a method for iteratively adjusting the weights (starting with zeros or random weights) so as to get as many training input-output pairs as possible right. However, hard thresholding entails that, for a given pair of an input pattern and a desired output pattern, small changes to the weights will often make no difference to the output. This makes it difficult to learn the weights for a multi-layer network by gradient descent, where small adjustments to the weights are made so as to iteratively reduce the errors. If the hard threshold is replaced by a soft threshold that varies continuously, such as a sigmoid function, gradient descent can be used for learning.

##### What is meant by “neural network”?

The term “neural network” originally refers to a network of biological neurons. More broadly, the term evokes a particular paradigm for understanding brain function, in which neurons are the essential computational units and computation is explained in terms of network interactions. Note that this leaves aside many biological complexities, including functional contributions of neurochemical diffusion processes, glia cells, and hemodynamics (Moore & Cao 2008). Although neurons are biological entities, the term “neural network” has come to be used as a shorthand for *artificial neural network,* a class of models of parallel information processing that is inspired by biological neural networks, but commits to several further major simplifications. Although spiking models have an important place in the computational literature, the models discussed here are nonspiking and do not capture dendritic computation, other processes within each neuron (e.g. Gallistel & King 2011), and distinct contributions from different types of neuron. The spatial structure of a neuron is typically abstracted from and its spiking output modelled as a real number analogous to the spike rate. The rate is modelled as a weighted sum of incoming “activations” passed through a static nonlinearity. Despite and because of these simplifications, the neural network paradigm provides one of the most important paths toward understanding brain information processing. It appears likely that this approach will take a central role in any comprehensive future brain theory. Opinions diverge as to whether more biologically detailed models will ultimately be needed. However, neural networks as used in engineering are certainly neurobiologically plausible and their success in artificial intelligence suggests that their abstractions may be desirable, enabling us to explain at least some complex feats of brain information processing.

### Networks with nonlinear hidden units are universal function approximators

The particular shape of the nonlinear activation function does not matter to the class of input-output functions that can be represented. Feedforward networks with at least one layer of “hidden” units intervening between input and output layers are universal function approximators, in the sense that, given a sufficient number of hidden units, they can approximate any function of the inputs in the output units. Continuous functions can be approximated with arbitrary precision by adding a sufficient number of hidden units and suitably setting the weights (Schäfer & Zimmermann 2007; Hornik 1991; Cybenko 1989). Fig. 2c illustrates this for two-dimensional inputs: Adding up a sufficient number of sigmoid ramps, which can have any orientation, slope, and position, we can approximate any continuous function of the input.

To gain an intuition on why combining sigmoids (or any step-like functions) of the input enables a network to approximate any function, imagine we set the weights of a set of units so that they compute sigmoids whose plateaus (close to 1) overlap only in a particular region of the input space. If we now sum the outputs of these units in a unit with a high threshold, that unit can indicate (by an output close to 1) that we are in a certain region of the input space. If we build indicators in this fashion for all regions within the input space that require a different output, we can map any input to any required output approximately. The precision of this approximate mapping can always be improved by using more units to define more separate regions with indicators. Note that if the activation function is continuous (as it usually is), then the function represented by the network is also continuous. The network would use two hidden layers to represent what is essentially a lookup table of the training input-output pairs. (However, it would have the nice feature of interpolating for novel intermediate inputs.)

The universal approximation theorem (Schäfer & Zimmermann 2007; Hornik 1991; Cybenko 1989) of feedforward nets goes beyond this intuitive explanation and shows that a single hidden layer is sufficient to approximate any continuous function and that the activation function need not resemble a step function.

A feedforward net is composed of a sequence of layers of units, with each unit sending its output only to units in higher layers (Fig 1b, c). The units and connections of a feedforward net, thus, correspond to the nodes and edges, respectively, of a directed acyclic graph. In computer vision systems, units often receive inputs only from the immediately preceding layer and inputs in lower layers are usually restricted to local receptive fields, inspired by the visual hierarchy.

Modern models use a variety of nonlinear activation functions, including sigmoid (e.g. logistic or hyperbolic tangent) and rectified linear ones (Fig. 1a). A rectified linear unit outputs the linear combination it computes, if it is positive, and 0 otherwise. Rectified linear units simplify the gradient-descent learning of the weights, enabling more rapid training, and work very well in computer vision and other domains.

### Why deep?

A feedforward net is called *deep* when it has more than one hidden layer. This technical definition notwithstanding, the term “deep” is also used in a graded sense. A deep net, thus, is a net with many layers and one net can be deeper than another. *Deep learning* refers to the strategy of using architectures with many hidden layers to tackle difficult problems, including vision.

This raises the question of why depth matters. We saw above that even shallow networks with a single layer of nonlinear hidden units are universal function approximators. Shallow networks are closely related to support vector machines, which can likewise learn arbitrary nonlinear functions, can be more efficiently trained than neural networks, and have been very successful tools of machine learning.

The reason depth matters is that deep nets can represent many complex functions more concisely (i.e. with fewer units and weights) than shallow nets and support vector machines (Bengio 2009).

Consider a shallow network (i.e. a net with a single hidden layer) that computes some function. We can create a deeper network with the same number of units by distributing the units of the single hidden layer across multiple hidden layers in the new net. The deep net could have the same connectivity from the input to the hidden units and from the hidden units to the output. It can thus compute any function the shallow net can compute. However, the reverse is not true. The deep net is permitted additional non-zero weights from any given layer to higher layers. This enables reuse of the results of previous computations and extends the deep net’s expressive power. For many possible functions that the deep net might compute, it can be shown that a shallow net would need a much larger number of units (Bengio 2009).

It is instructive to consider the analogy to modern computing hardware. The von Neumann architecture is a fundamentally sequential model of computation that enables the reuse of results of previous computations. In special cases, where many computations can be performed independently, parallel hardware can speed up the process. However, whereas independent computations can be performed either in parallel or sequentially, dependent computations can *only* be performed sequentially. The option to reuse previous results therefore extends the set of computable functions (if the total number of units is fixed).

In essence, a shallow network is a universal function approximator because it can piece together the target function like a lookup table. However, many functions can be more concisely represented using a deeper net, taking advantage of redundancies and exploiting the inherent structure of the target function. Although every problem is different and the field is still learning when exactly depth helps, the practical success of deep learning in AI suggests that many real-world problems, including vision, may be more efficiently solved with deep architectures. Interestingly, the visual hierarchy of biological brains is also a deep architecture.

### Recurrent neural networks are universal approximators of dynamical systems

Feedforward nets compute static functions. An architecture with more interesting dynamics is a recurrent net, whose units can be connected in cycles. This is more similar to biological neuronal networks, in which lateral and feedback connections are ubiquitous. Note that the notion of separate hidden layers is meaningless in a recurrent net because each hidden unit can interact with each other hidden unit. Recurrent nets are therefore often depicted as a single interconnected set of hidden units, with separate sets of input and output units (Fig. 1d). A layered architecture is a special case of a recurrent network in which certain connections are missing (i.e. their weights fixed at 0).

In visual neuroscience, the theoretical concept of the visual hierarchy is based on a division of the connections into feedforward, lateral, and feedback, and on the fact that some neurons are separated from the input by a larger number of synapses and tend to represent more complex features of the input. Although these criteria may not support a perfectly unambiguous assignment of ranks that would define a hierarchy for the primate visual system (Hilgetag et al. 2000), the hierarchical model continues to be a useful simplification.

Whereas a feedfoward network computes a static function mapping inputs to outputs, a recurrent network produces dynamics: a temporal evolution of states that can be influenced by a temporal evolution of input patterns. The internal state of a recurrent net lends it a memory and enables it to dynamically compress the recent stimulus history and efficiently detect temporal patterns. Whereas feedforward nets are universal function approximators, recurrent nets are universal approximators of dynamical systems (Schäfer & Zimmermann 2007). A variety of particular models have been explored by simulation and analytically.

In an echo-state network (Jaeger 2001, see also Maass et al. 2002, for a similar model with spiking dynamics), for example, the sequence of input patterns is fed into a set of hidden units that are sparsely and randomly connected. The wave of activity associated with each input pattern will reverberate among the hidden units for a while until it comes to be dominated by the effects of subsequent input patterns. Like concentric waves on the surface of a pond that enable us to infer an event at their center sometime in the past, the activity of the hidden units encodes information about the recent stimulus history. In echo-state networks, the weights among the hidden units are not trained (although their random setting requires some care to ensure that the memories do not fade too quickly). Supervised learning is used to train a set of readout units to detect temporal patterns in the input.

Echo-state networks rely on random weights among the hidden units for their recurrent dynamics. Alternatively, the dynamics of a recurrent net can be explicitly learned through supervision, so as to optimise a recurrent net to produce, classify, or predict certain temporal patterns (Graves & Schmidhuber 2009, Sutskever et al. 2014).

### Representations can be learned by gradient descent using the backpropagation algorithm

The universality theorems assure us of the representational power of neural networks with sufficient numbers of units. However, they do not tell us how to set the weights of the connections, so as to represent a particular function with a feedforward net, or a particular dynamical system with a recurrent net. Learning poses a high-dimensional and difficult optimisation problem. Models that can solve real-world problems will have large numbers of units and even larger numbers of weights. Global optimisation techniques are not available for this nonconvex problem. The space of weight settings is so vast that simple, e.g. evolutionary, search algorithms can only cover a vanishingly small subset of the possibilities and typically do not yield working solutions, except for small models restricted to toy problems.

The high dimensionality of the weight space makes global optimisation intractible. However, the space contains many equivalent solutions (consider, for example, exchanging all incoming and outgoing weights between two neurons). Moreover, the total error (i.e., the sum of squared deviations between actual and desired outputs) is a locally smooth function of the weights.

The current training method of choice is gradient descent, the iterative reduction of the errors through small adjustments to the weights.

The basic idea of gradient-descent learning is to start with a random initialisation of the weights and to determine, for each weight, how much a slight change to it will reduce the error. The weight is then adjusted in proportion to the effect on the error. This method ensures that we move in the direction in weight space, along which the error descends most steeply.

The gradient, i.e. how much the error changes with an adjustment of a weight, is the derivative of the error with respect to the weight. These derivatives can be computed easily for the weights connecting to the output layer of a feedforward net. For connections driving the preceding layers, an efficient way to compute the error derivatives is to propagate them backward through the network. This gives the method its name *backpropagation* (Werbos 1981; Rumelhart et al. 1986).

Gradient descent sees only the local neighborhood in weight space and is not guaranteed to find globally optimal solutions. It can nevertheless find solutions that work very well in practice. The high dimensionality of the weight space is a curse in that it makes global optimisation difficult. However, it is a blessing in that it helps local optimisation find good solutions: with so many directions to move in, gradient descent is unlikely to get stuck in local minima, where the error increases in all directions and no further progress is possible.

Intriguingly, the same approach can be used to train recurrent networks. The error derivatives are then computed by *backpropagation through time.* To understand this we can construe a recurrent network as the feedforward network obtained by replicating the recurrent net along the dimension of time (for a sufficiently large number of time steps). A recurrent net is equivalent to a feedforward net with an infinite number of layers, each connected to the next by the same weights matrix (the recurrent net’s weights matrix).

By backpropagation through time, a recurrent net can learn weights that enable it to store short-term memories in its dynamics and relate temporally separated events, so as to achieve the desired classifications or predictions. However, propagating error derivatives backward through time far enough for the net to learn to exploit long-lag dependencies is hampered by the problem that the gradients tend to vanish or explode (Hochreiter 1991; Hochreiter et al. 2001). The problem occurs because a given weight’s error derivative is the product of multiple terms, corresponding to weights and derivatives of the activation functions encountered along the path of backpropagation. One solution to this problem is offered by the long short-term memory (LSTM) architecture (Hochreiter & Schmidhuber, 1997), in which special gated units can store short-term memories for extended periods. The error derivatives backpropagated through these units remain stable, enabling backpropagation to learn long-lag dependencies. Such networks, amazingly, can *learn to remember* information that will be relevant many time steps later in a sequential prediction or control task. Backpropagation adjusts the weights to ingrain a dynamics that selectively stores (in the activation state) information needed later to perform the task.

Vanishing and exploding gradients also pose a problem in training deep feedforward nets with backpropagation. The problem occurs because a given weight’s error derivative is the product of weights and derivatives of the activation function encountered along the path of backpropagation. The choice of nonlinear activation function can make a difference with regard to this problem. In addition, the details of the gradient descent algorithm, regularisation, and weight initialisation all matter to making supervised learning by backpropagation work well.

### Representations can also be learned with unsupervised techniques

In supervised learning, the training data comprise both input patterns and the associated desired outputs. An explicit supervision signal of this type is often unavailable in the real world. Biological organisms do not in general have access to explicit supervision. In engineering, similarly, we often have a large number of unlabeled and only a smaller number of labeled input patterns (e.g. images from the web). Unsupervised learning does not require labels for a network to learn a representation that is optimised for natural input patterns and potentially useful for a variety of tasks. Natural images, for example, form a very small subset of all possible images. This enables unsupervised learning to find compressed representations for natural images.

An instructive example of unsupervised learning is provided by *autoencoders* (Hinton & Salakhutdinov 2006). An autoencoder is a feedforward neural network with a central “code layer” that has fewer units than the input. The network is trained with backpropagation to reconstruct its input in the output layer (which matches the input layer in its number of units). Although the learning algorithm is backpropagation and uses a supervision signal, the technique is unsupervised because it requires no separate supervision information (i.e. no labels) but only the set of input patterns. If all layers, including the code layer, had the same dimensionality as the input, the net could just pass the input through its layers. Since the code layer has fewer units, however, it forms an informational bottleneck. In order to reconstruct the input, the net must learn to retain sufficient information about the input in its small code layer. An autoencoder therefore learns a compressed representation in its code layer. This representation will be specialised for the distribution of input patterns that the autoencoder has been trained with.

The layers from the input to the code layer are called the *encoder.* The layers from the code layer to the output are called the *decoder.* If encoder and decoder are linear, the network learns the linear subspace spanned by the first k principal components (for a code layer of k units). With nonlinear neural nets as encoders and decoders, nonlinear compressed representations can be learned. Nonlinear codes can be substantially more efficient when the natural distribution of the input patterns is not well represented by a linear subspace. Natural images are a case in point.

Unsupervised learning can help pretrain a feedforward network when insufficient labeled training data are available for purely supervised learning. For example, a net for visual recognition can be pretrained layer by layer in the autoencoder framework, using a large set of unlabeled images. Once the net has learned a reasonable representation of natural images, it can more easily be trained with backpropagation to predict the right image labels.

## FEEDFORWARD NEURAL NETWORKS FOR VISUAL OBJECT RECOGNITION

Computer vision has recently come to be dominated by a particular type of deep neural network: the deep feedforward convolutional net. These networks now robustly outperform the previous state of the art, which consisted in hand-engineered visual features (e.g. Lowe 1999) forming the input to shallow machine learning classifiers, such as support vector machines. Interestingly, some of the earlier systems inserted an intermediate representation, often acquired by unsupervised learning, between the hand-engineered features and the supervised classifier. This might have helped address the need for a deeper architecture.

The deep convolutional nets dominating computer vision today share a number of architectural features, some of which are loosely inspired by biological vision systems (Hubel & Wiesel 1968).

- **Deep hierarchy**: These networks typically have 5 to 10 layers, comparable to the number of stages along the primate ventral visual stream (Fig. 3). They process information through a hierarchy of representations, gradually transforming a visual representation, whose spatial layout matches the image, to a semantic representation that enables the recognition of object categories.
- **Convolution**: Early layers contain local feature detectors. Each detector is replicated all over the two dimensional image, forming a feature map. This amounts to a convolution of the image with each local feature pattern, followed by a static nonlinearity. The convolutional architecture is motivated by the insight that a feature that is useful in one position is likely to also be useful in another position and resembles biological vision, where local features are also replicated across the visual field (though not quite uniformly as in convolutional nets). The resulting receptive fields (RFs) of the units are localised in the visual field and increase in size along the hierarchy. The restriction of the connections to a local region and the replicati.on of the connection weights across spatial positions (same weight pattern at all locations for a given feature) greatly reduces the number of independent weights that need to be learned (LeCun 1989).
- **Local pooling and subsampling**: In between the convolutional layers, local pooling layers are inserted. Pooling combines the outputs of a local set of units in the previous layers by taking the maximum or the average. This confers a local tolerance to spatial shifts of the features, making the representation robust to small image shifts and small distortions of the configuration of image features (Fukushima 1980). Max-pooling is also used in neuroscientific vision models like HMAX (Riesenhuber & Poggio 1999; Serre et al. 2007) to implement local tolerances. Pooling is often combined with subsampling of the spatial locations. The reduction in the number of distinct spatial locations represented frees up resources for an increase along the hierarchy in the number of distinct features computed at each location.

**Figure 3:**
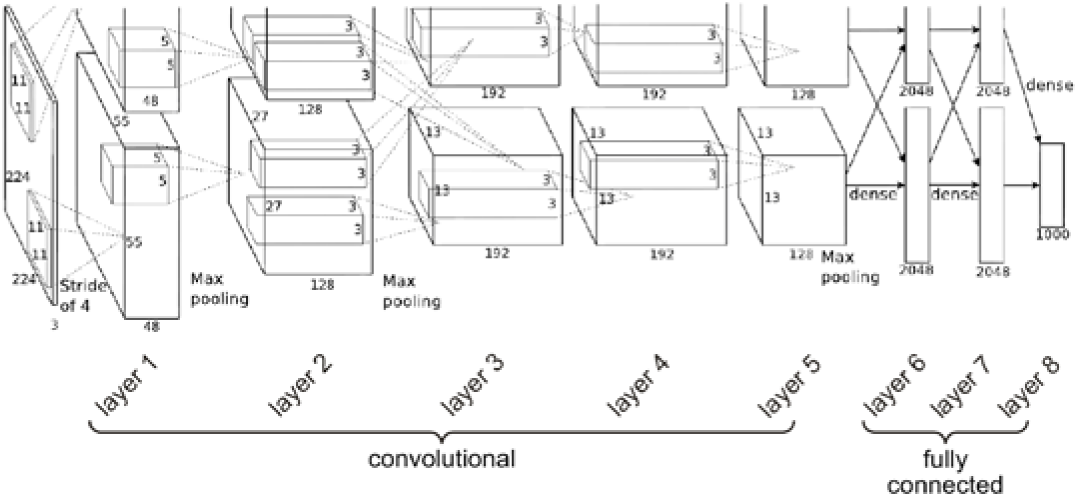
Deep convolutional feedforward architecture for object recogition. The figure shows the architecture used by Krizhesvsky et al. (2012). The information flows from the input pixel image (left, 224 pixels by 224 pixels, 3 colour channels) through 7 hidden layers to the category output (right, 1000 category detector units). The large boxes represent stacks of feature maps. For layer 2, for example, the lower large box represents 128 feature maps of size 27 (horizontal image positions) by 27 (vertical image positions). Note that the dimensions of the boxes are not drawn to scale. The small boxes represent the feature templates that are convolved with a given layer’s representation. Because convolution and maxpooling operate at strides greater than 1 pixel, the spatial extent of the feature maps decreases along the sequence of representations (224, 55, 27, 13, 13, 13, 1, 1, 1). Upper and lower large boxes represent the division of labour between 2 graphics processing units.

In the highest layers, units have global RFs, receiving inputs from all units of the previous layer. The final layer typically contains one unit per category and implements a softmax (normalised exponential), which strongly reduces all but the very highest responses and ensures that the outputs add up to 1. When the crossentropy error is used in training, the output can be interpreted as a probability distribution over the categories.

The networks can be trained to recognize the category of the input image using backpropagation (LeCun et al. 1989, LeCun & Bengio 1995). When a network is trained to categorize natural images, the learning process discovers features that are qualitatively similar to those found in biological visual systems (Figure 4). The early layers develop Gabor-like features, similar to those that characterize V1 neurons. Similar features are discovered by unsupervised techniques such as sparse representational learning (Olshausen & Field 1997), suggesting that they provide a good starting point for vision, whether the goal is sparse representation or categorization. Subsequent stages contain units that are selective for slightly more complex features, including curve segments. Higher layers contain units that are selective for parts of objects and for entire objects, such as faces and bodies of humans and animals, and inanimate objects such as cars and buildings.

**Figure 4:**
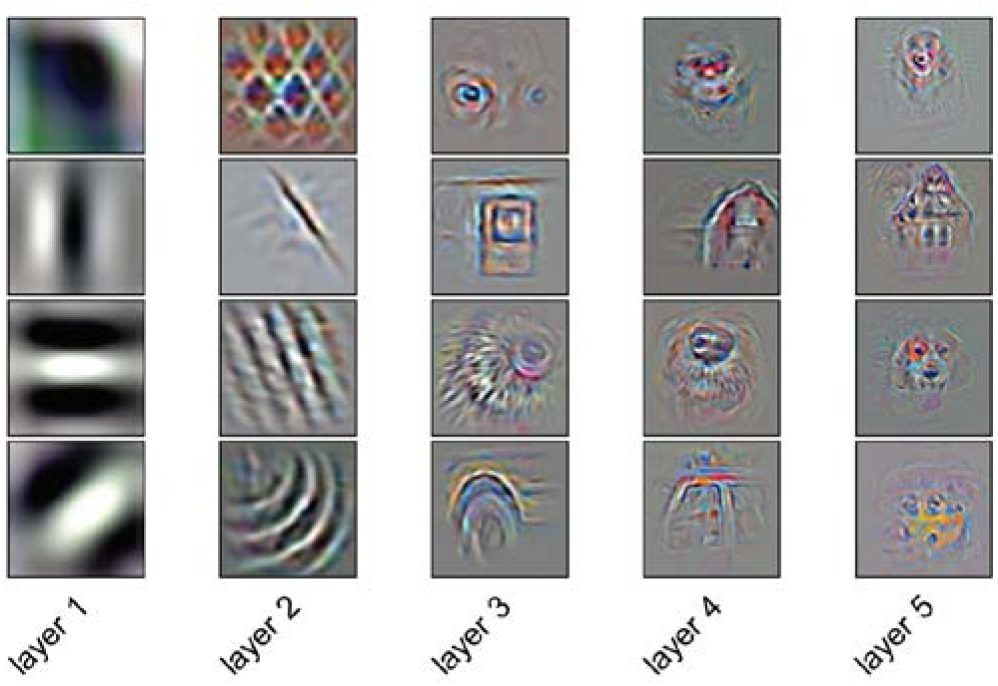
Deep supervised learning produces feature selectivities qualitatively consistent with neuroscientific theory. In order to understand representations in deep neural networks, we can visualise what image elements drive a given unit in a deep net. For 20 example units (4 units from each of the layers), the images shown visualise what caused the response in the context of a particular image that strongly drove the unit. The visualisation technique used here involves two steps. First an input image strongly driving the unit is selected. Then the feedforward computation driving the unit is inverted to generate the image element responsible. Convolutions along the feedforward pass are inverted by deconvolution (using the transposes of the convolution matrices). Maxpooling operations are inverted by storing which connection to the pooling unit was maximally active in the feedforward pass. Note that a unit deep in a network does not perform a simple template matching operation on the image and therefore cannot be fully characterised by any visual template. However, performing the above visualisation for many images that strongly drive a unit (not shown) can help us understand a unit’s selectivity and tolerances. The deconvolutional visualisation technique shown was developed by Zeiler & Fergus (2013). The deep net is from Chatfield et al. (2014). The analysis was performed by Güçlü & van Gerven (2015). Figure adapted from Güçlü & van Gerven (2015).

To understand what has been learned automatically, the field is beginning to devise methods for visualising units’ receptive fields and selectivities (Zeiler & Fergus 2013; Girshick et al. 2014; Simonyan et al. 2014; Tsai & Cox 2015; Zhou et al. 2015). Fig. 4 shows such visualisations, which support the idea that units learn selectivities to natural image features whose visual complexity increases along the hierarchy. However, there are two important caveats to such visualisations: First, because of the multiple nonlinear transforms across layers, a unit cannot be accurately characterised by an image template. If the high-level responses could be computed by template matching, there would be no need for a deep hierarchy. The visualisations merely show what drives the response in the context of a particular image. Many images driving a unit need to be considered to get an idea of its selectivity (for multiple templates for each of a larger number of units, see Zeiler & Fergus 2013). Second, the units visualised in Fig. 4 have been selected because they confirm a theoretical bias for interpretable selectivities. Units like those shown may be the exception rather than the rule, and it is unclear whether they are essential to the network’s functionality. For example, meaningful selectivities could reside in linear combinations of units rather than single units, with weak distributed activities encoding essential information.

The representational hierarchy appears to gradually transform a space-based visual to a shape-based and semantic representation. The network acquires complex knowledge about the kinds of shapes associated with each category. In this context, shape refers to luminance- and color-defined features of various levels of complexity. High-level units appear to learn representations of shapes occurring in natural images, such as faces, human bodies, animals, natural scenes, buildings, and cars. The selectivities learned are not restricted to the categories detected by the output layer, but may include selectivities to parts of these objects or even to context elements. For example, the network by Krizhesvsky et al. (2012) contains units that appear to be selective for text (Yosinski et al. 2015) and faces, although text and faces were not among the trained categories. Presumably, those responses help detect the categories represented in the output layer, because they are statistically related to the categories to be detected. For example, part-selective features may serve as stepping stones toward detection of entire objects (Jozwik et al. 2015). A verbal functional interpretation of a unit, e.g. as an eye or a face detector, may help our intuitive understanding and capture something important. However, such verbal interpretations may overstate the degree of categoricality and localisation, and understate the statistical and distributed nature of these representations.

An influential example of a deep convolutional neural net for computer vision is the system built by Krizhesvsky et al. (2012). The architecture (Fig 3) comprises five convolutional and three fully connected layers. The authors found that reducing the number of convolutional layers hurt performance, illustrating the need for a deep architecture. The system uses rectified linear units, max-pooling, and local normalisation. The network was trained by backpropagation to recognise which of 1,000 object categories was shown in the input image. The training set comprised 1.2 million category-labelled images from the ImageNet set. This set was expanded by a factor of 2048, by adding translated and horizontally reflected versions of the images. The training cycled 90 times through the resulting image set.

The training relied on dropout regularisation (Hinton et al. 2012), where each unit is “dropped” with a probability of 0.5 on each trial. On a given learning trial, thus, a random set of about half of the units is used in both the forward pass computing the output and the backpropagation pass adjusting the weights. This method prevents complex co-adaptations of the units during learning, forcing each unit to make a useful contribution in the context of many different “teams” of other units. The network has a total of 650,000 units and 60 million parameters. The convolutional layers are defined by their small local weight templates, which constitute less than 5% of the parameters in total. Over 95% of the parameters define the upper three fully connected layers. Dropout was applied to the first two fully connected layers, each of which has many millions of incoming connections. Experiments showed that dropout was necessary to prevent overfitting.

The training was performed over the course of six days on a single workstation with two graphics processing units (GPUs), which parallelise and greatly accelerate the computations. The system was tested on a held-out set of images in the ImageNet Large-Scale Visual Recognition Challenge 2012, a computer vision competition. It won the competition, beating the second best system by a large margin and marking the beginning of the dominance of neural nets in computer vision. Since then a number of convolutional neural networks using similar architectures have further improved upon its performance (e.g. Zeiler & Fergus 2013; Chatfield et al. 2014).

The deep convolutional neural nets of computer vision perform limited aspects of vision, such as category-level recognition. However, the range of visual tasks tackled is quickly expanding and deep nets do represent a quantum leap compared to the earlier computer vision systems. Deep convolutional nets are not designed to closely parallel biological vision systems. However, their essential functional mechanisms are inspired by biological brains and could plausibly be implemented with biological neurons. This new technology provides an exciting framework for more biologically faithful brain-computational models that perform complex feats of intelligence beyond the current reach of computational neuroscience.

#### Adversarial examples can reveal idiosyncrasies of neural network architectures

Fooling vision can help us learn about its mechanisms. This goes for biological as well as artificial vision. Researchers are exploring how artificial neural nets represent images by trying to fool them (Szegedy et al. 2014; Nguyen et al. 2015; Goodfellow et al. 2015). They use optimisation techniques to design images that are incorrectly classified. An adversarial example is an image from category X (e.g. a bus or a noise image) that has been designed to be misclassified by a particular network as belonging to category Y (e.g. an ostrich). This can be achieved by taking a natural image from category X and adjusting the pixels so as to fool the net. The backpropagation algorithm, which usually serves to find small adjustments to the weights that reduce the error for an image, can be used to find small adjustments to the image that *create* an error. For the convolutional neural nets currently used in computer vision, adversarial examples can be created by very slight changes to the image that clearly do not render it a valid example of a different category. An adversarial example can look indistinguishable from the original image to humans. This has been taken as evidence of the limitations of current neural net architectures as vision systems and as models of human vision.

An adversarial example created to fool an artificial neural net will not usually fool a human observer. However, it is not known whether adversarial examples can be created for human visual systems as well. Constructing adversarial examples requires computational optimisation based on full knowledge of the particular network to be fooled. We cannot currently match this process with psychophysical and neurophysiological techniques to fool biological vision. An intriguing possibility, thus, is that biological visual systems, too, are susceptible to adversarial examples.

These could exploit idiosyncrasies of a particular brain, such that an adversarial example created to fool one person will not fool another person. The purpose of vision systems is to work well under natural conditions, not to be immune to extremely savvy sabotage that requires omniscient access to a network’s internal structure and precise stabilisation of the fooling image on the system’s retina (or visual sensor array). Human vision is famously susceptible to visual illusions of various kinds. From a machine learning perspective, it appears inevitable that adversarial examples can be constructed for any learning system – artificial or natural – that must rely on an imperfect inductive bias to generalise from a limited set of examples to a high-dimensional classification function.

What lessons do the adversarial examples hold about current neural net models? If an adversarial example fooled only the particular instance of a network it was constructed for, exploiting that particular net’s idiosyncrasies, then they would be easy to dismiss. However, adversarial examples generalise across networks to some extent. If a new network is created by initializing the same architecture with new random weights and training it with the same set of labeled images, the resulting network will often still be fooled by an adversarial example created for the original network. Adversarial examples also generalize to slightly altered architectures if the same training set is used. If the training set is changed, adversarial examples created for the original network are not very effective anymore, but they may still be misclassified at a higher rate than natural images. This suggests that adversarial examples exploit network idiosyncrasies resulting largely from the training set, but also to some extent from the basic computational operations used. One possibility is that the linear combination computed by each unit in current systems makes them particularly easy to fool (Goodfellow et al. 2015). In essence, each unit divides its input space by a linear boundary (even if its activation rises smoothly as we cross the boundary for sigmoid or linearly on the preferred side for rectified linear activation functions). By contrast, networks using radial basis functions, where each unit has a particular preferred pattern in its input space and its response falls off in all directions, might be harder to fool. However, they are also harder to train – and perhaps for the same reason. It will be intriguing to see this puzzle solved as we begin to compare the complex representational transformations between artificial and biological neural nets in greater detail.

## EARLY STUDIES TESTING DEEP NEURAL NETS AS MODELS OF BIOLOGICAL BRAIN REPRESENTATIONS

Several studies have begun to assess deep convolutional neural nets as models for biological vision, comparing both the internal representational spaces and performance levels between models and brains. One finding replicated and generalised across several studies (Yamins et al. 2014; Khaligh-Razavi & Kriegeskorte 2014) is that models that perform better at object recognition utilise representational spaces that are more similar to those of inferior temporal (IT) cortex (Tanaka 1996) in human and nonhuman primates. This affirms the intuition that engineering approaches to computer vision can learn from biological vision.

It is not true in general that engineering solutions closely follow biological solutions (consider planes, trains, and automobiles). In computer vision, in particular, early failures to scale neural net models to real-world vision fostered a sense that seeking more brain-like solutions was fruitless. The recent successes of neural net models suggest that brain-inspired architectures for vision are extremely powerful. The empirical comparisons between representations in computer vision systems and brains discussed in this section further suggest that neural net models learn representations very similar to those of the primate ventral visual pathway. Although it is impossible to prove that representations have to be similar to biological brains to support successful computer vision, several studies comparing many models reported that representations more similar to IT perform better at object recognition (Yamins et al. 2014, Khaligh-Razavi et al. 2014).

The association between performance and representational similarity to IT has been shown for a large set of automatically generated neural net architectures using random features (Yamins et al. 2013; 2014), for a wide range of popular hand-engineered computer vision features and neuroscientific vision models (Khaligh-Razavi et al. 2013; 2014), and for the layers of a single deep neural network (Khaligh-Razavi et al. 2014). At least within the architectures explored so far, it appears that performance optimisation leads to representational spaces similar to IT.

IT is known to emphasise categorical divisions in its representation (Kriegeskorte et al. 2008b). Models that perform well at categorisation (which is implemented by linear readout) likewise tend to have stronger categorical divisions. This partially explains their greater representational similarity to IT. However, even the within-category representational geometries tend to be more similar to IT in the better performing models (Khaligh-Razavi et al. 2014).

The best performing models are deep neural nets and they are also best at explaining the IT representational geometry (Khaligh-Razavi et al. 2014; Cadieu et al. 2014). Khaligh-Razavi et al. (2014) tested a wide range of classical computer vision features, several neuroscientifically motivated vision models, including VisNet (Wallis & Rolls 1997; Tromans et al. 2011) and HMAX (Riesenhuber & Poggio 1999), and the deep neural network of Krizhesvsky et al. (2012; Fig. 3). The brain representations were estimated from human functional magnetic resonance imaging (fMRI) and monkey cell recordings (data from Kriegeskorte et al. 2008b; Kiani et al. 2007).

They compared the internal representational spaces between models and brain regions using representational similarity analysis (Kriegeskorte et al. 2008a). For each pair of stimuli, the dissimilarity of the two stimuli in the representation is measured. The vector of representational dissimilarities across all stimulus pairs can then be compared between a model representation and a brain region.

Early layers of the deep neural net had representations resembling early visual cortex. Across the layers of the net, the representational geometry became monotonically less similar to early visual cortex and more similar to IT. These results are shown in Fig. 5 for human data; IT results were similar for human and monkey.

**Figure 5:**
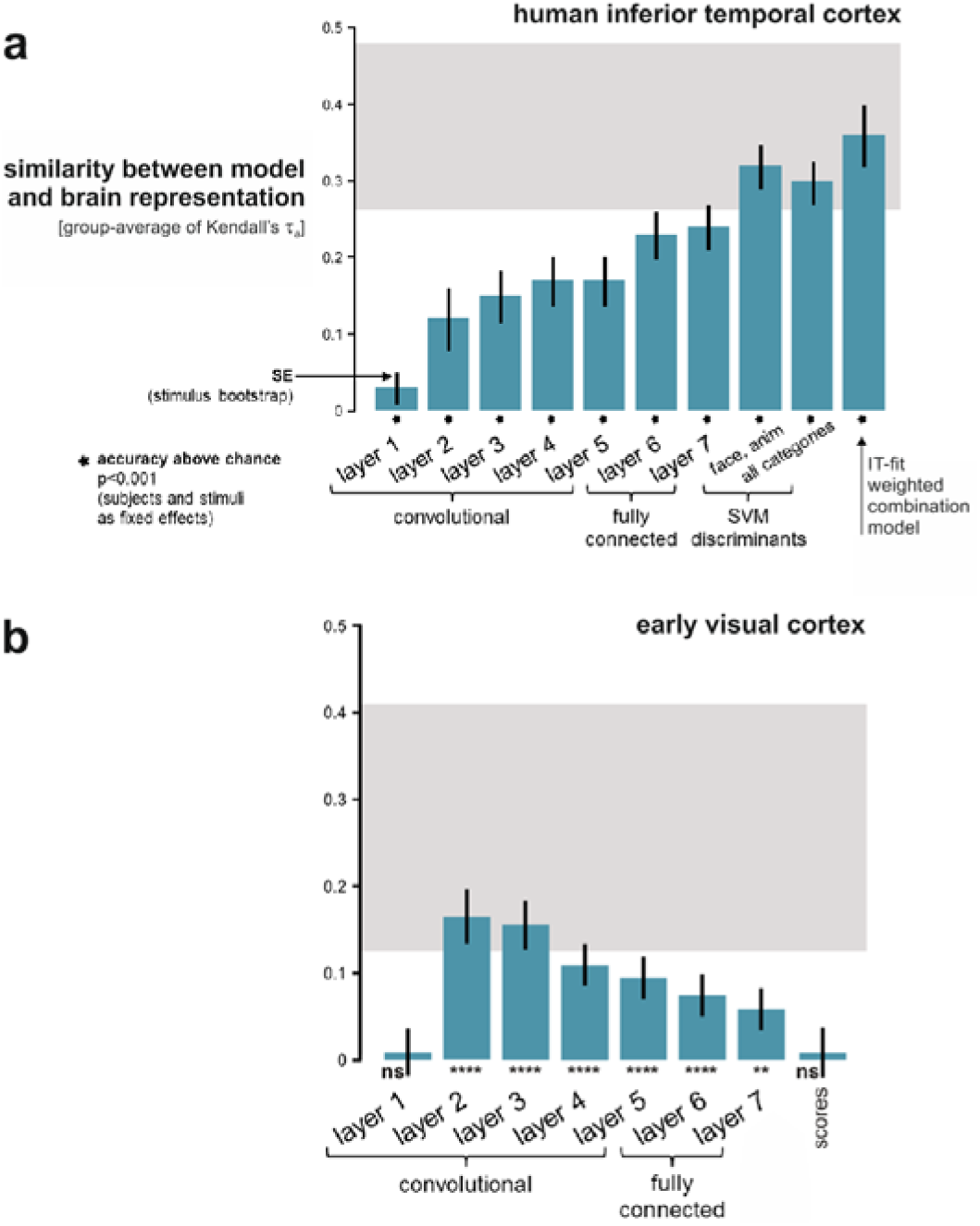
Deep neural net explains early visual and inferior temporal representations of object images. Representations in model and brain were characterised by the dissimilarity matrix of the response patterns elicited by a set of real-world object images. (a) As we ascend the layers of the Krizhevsky et al. (2012) neural net, the representations become monotonically more similar to that of human inferior temporal (IT) cortex. When the final representational stages are linearly remixed to emphasise the same semantic dimensions as IT, the noise ceiling (gray) is reached. Similar results obtain for monkey IT. (b) Lower layers of the deep neural net resemble the representations in the foveal confluence of early visual areas (V1-3). Results reproduced from Khaligh-Razavi & Kriegeskorte (2014).

At the highest layer, the representation did not yet fully explain the explainable (non-noise) variance in the IT data. However, a representation fitted to IT by linear remixing and reweighting of the deep neural net’s features (using independent image sets for training and validation), fully explained the IT data (Khaligh-Razavi et al. 2014). This IT-fitted deep neural net representation explained the IT representation substantially and significantly better than a similarly IT-fitted combination of the conventional computer vision features.

Cadieu et al. (2013; 2014) analysed the internal representations of a population of IT cells alongside models of early vision, the HMAX model, a hierarchically optimised multi-layer model from Yamins et al. (2013; 2014), and the deep neural nets from Krizhevsky et al. (2012) and Zeiler & Fergus (2013). The representations performing best at object categorisation (Fig. 6a) were the deep net of Zeiler & Fergus (2013) and the biological IT representation (monkey neuronal recordings), followed closely by the deep net of Krizhevsky et al. (2012). The other representations performed at much lower levels. The two deep nets explained the IT data equally well as did neuronal recordings from an independent set of IT neurons (Fig. 6b).

**Figure 6:**
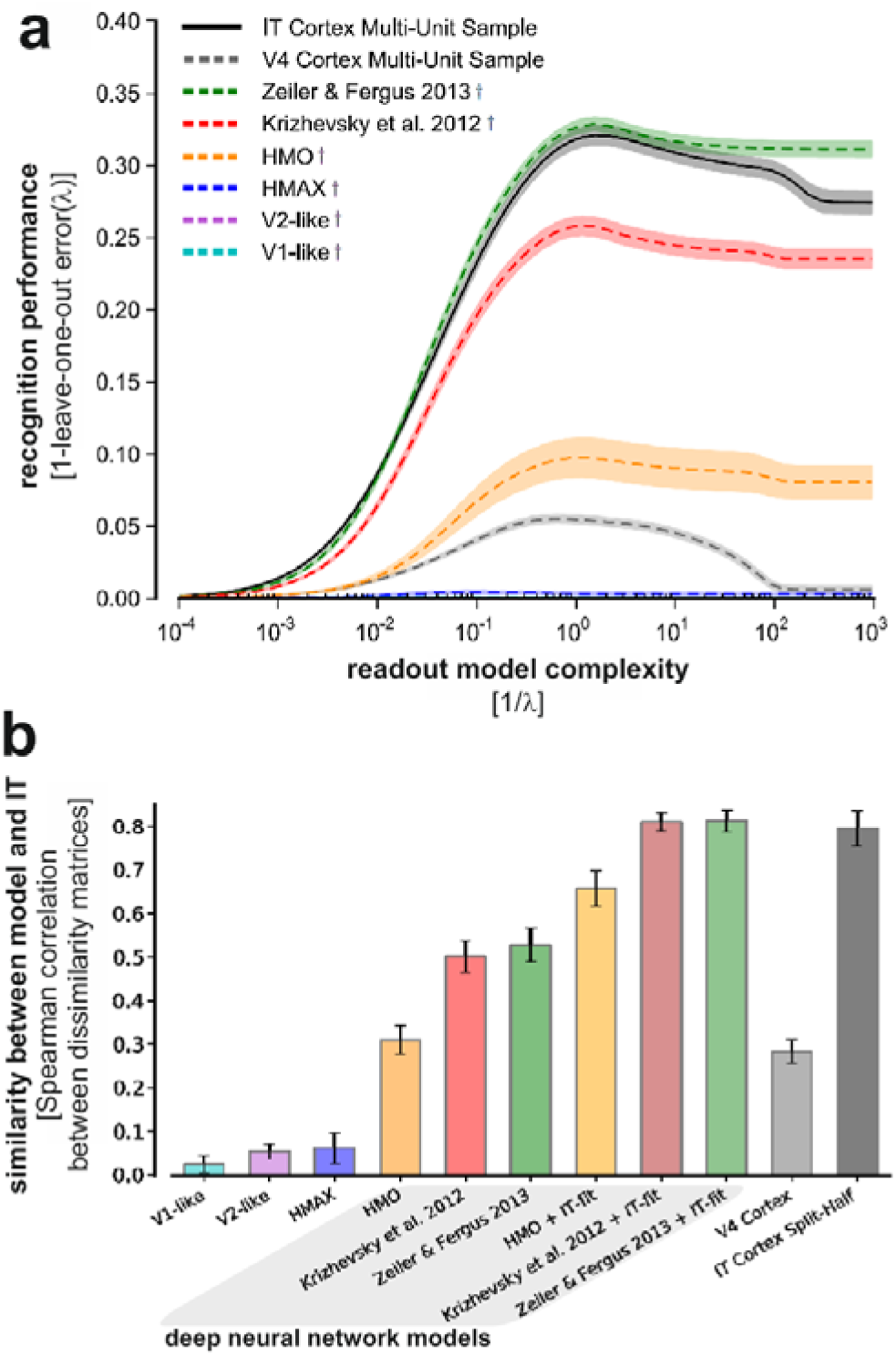
Deep neural networks beat simpler computational models at recognition and better explain IT representations. (a) Object recognition performance of deep neural networks beats that of shallower models and rivals that of a population of IT neurons recorded in a monkey. Recognition performance (vertical axis) is plotted as a function of readout-model complexity (horizontal axis); high performance at low complexity indicates that the categories occupy easily separable regions in the representational space. (b) Deep neural network representations more closely resemble IT than do three simpler models (V1-like, V2-like, and HMAX). The similarity between each model and IT (vertical axis) was measured using the Spearman’s rank correlation coefficient to compare representational dissimilarity matrices. Results reproduced from Cadieu et al. (2014). Abbreviations: HMAX, hierarchical model and X (Riesenhuber & Poggio 1999, Serre et al. 2007, Tarr 1999); HMO, hierarchical modular optimization model (Yamins et al. 2014); IT, inferior temporal cortex.

Several further studies have yielded similar results and are beginning to characterise to what extent representations at different depths can explain the representational stages of the ventral stream (Agrawal et al. 2014, Güçlü & van Gerven 2014, Khaligh-Razavi et al. 2015).

Overall these early attempts to empirically compare representations between deep neural net object recognition models and inferior temporal cortex suggest four conclusions: (1) Only deep neural nets perform object recognition at levels comparable to humans. (2) Only deep neural nets explain the representational geometry of IT. (3) The representation appears to be gradually transformed with lower layers resembling the earlier stages of the primate ventral stream. (4) The high-level deep net representation resembles IT not merely in that it emphasises categorical divisions, but also in its within-category representational geometry.

## RECURRENT NEURAL NETS FOR VISION

Feedforward nets are useful as models of the initial sweep of neuronal signalling through the visual hierarchy. They go some way toward explaining vision at a glance. However, feedforward nets are unlike the brain in terms of their connectivity and dynamics and fundamentally limited to the computation of static functions. Rather than computing a static function on each of a series of image frames, vision takes a time-continuous input stream and interprets it through ongoing recurrent computations. The current network state likely represents the recent stimulus history along with predictions of impending events and other behaviourally important information.

Recurrent computations probably contribute to the automatic rapid interpretation of even static visual images (Sugase et al. 1999, Brincat & Connor 2006, Freiwald & Tsao 2010, Carlson et al. 2013, Cichy et al. 2014; Tang et al. 2014a), leading to the emergence of representations that clearly distinguish particular objects and object categories. Individual faces, for example, become more clearly distinguishable in monkey neuronal population codes at latencies that exceed the 100 ms or so it takes the feedforward sweep to reach IT (Sugase et al. 1999; Freiwald & Tsao 2010). At the level of object categories, similarly, evidence from human magnetoencephalography suggests that strong categorical divisions arise only at latencies of over 200 ms after stimulus onset (Carlson et al. 2013, Cichy et al. 2014). Both category and exemplar representations, thus, may rely on recurrent processing to achieve invariance to irrelevant variation among images that carry the same essential meaning.

The brain might rely on a combination of feedforward and recurrent processing to arrive at a representation similar to that computed in feedforward convolutional nets trained for object recognition. We have seen that a recurrent network can be unfolded as a deep feedforward network. Conversely, when the numbers of units and connections are limited, a desirable function computed by a very large feedforward network might alternatively be approximated by recurrent computations in a smaller network. Recurrent dynamics can expand computational power by multiplying the limited physical resources for computation along time.

Recurrent neuronal dynamics likely also serve more sophisticated computations than those of feedforward convolutional networks. Assume a visual neuron’s function is to represent the presence of some piece of content in the image (a feature, an object part, an object). The feedforward sweep alone might not provide the full evidence the neuron needs to confidently detect the piece of content it represents. The neuron might therefore integrate later-arriving lateral and top-down signals to converge on its ultimate response. Multiple neurons might pass messages recurrently until the population converges on a stable interpretation of the image.

Recurrent computations might implement the iterative fitting to the image of a generative model of image formation, with the fitted parameters specifying the contents (and causes) of the image (see Sidebar *The deep mystery of vision: How to integrate generative and discriminative models*). Assume, for simplicity, that the generative model is exactly invertible. This might be plausible if the model includes prior world knowledge sufficient to disambiguate visual images. Images and symbolic descriptions of their contents are then related by a one-to-one (bijective) mapping. In principle, the inverse of the generative model could be represented by a feedforward model (because of universality). However, such a model might require too many neurons and connections, or its connectivity might be impossible to learn from limited training data. Instead of analysing the image through feedforward computations, we can perform analysis by synthesis (Yuille & Kersten 2006), fitting a generative model of image formation to the particular image to be recognised.

The inversion of generative models has long been explored in both brain science and computer vision (Knill et al. 1996; Yuille & Kersten 2006; Prince 2012). The inference problem is hard because a vast number of combinations of surfaces and lights can explain any image. In order to constrain the search space and disambiguate the solution, the brain must use massive prior knowledge about the world. Inference on a generative model might be tractible if it were performed on a higher-level representation of the image computed by discriminative mechanisms. How the brain combines discriminative computations with inference on generative models to perceive the world is one of the fundamental unsolved problems of brain science.

The Helmholtz machine (Dayan et al. 1995) uses analysis by synthesis at the level of learning. A bottom-up recognition model and a top-down generative model are concurrently learned so as to best represent the distribution of the inputs in a maximum-likelihood sense. The learning can be performed using the wake-sleep algorithm (Hinton et al. 1995; Dayan 2000). In the wake phase, the recognition model “perceives” training images and the generative model learns to better reconstruct these images from their internal representations. In the sleep phase, the generative model “dreams” of images and the recognition model learns to better infer the internal representations from the images. By alternating wake and sleep phases, the two models co-adapt and jointly discover a good representation of the probability distribution over images.

A recurrent network could use the feedforward sweep to compute an initial rough estimate of the causes and subsequent recurrent computations to iteratively reduce the prediction error of the generative model and explain nonlinear interactions of the parts, such as occlusion. The process could use predictive coding (Lee & Mumford 2003; Friston 2010), with recognised parts of the image explained away (and subtracted out of) lower-level representations, such that the parts yet unexplained are gradually uncluttered in the low-level representation and contextualised in the high-level representation as the easier and then the harder bits of the image are successively recognised.

#### The deep mystery of vision: How to integrate generative and discriminative models

The current advances in computer vision are largely driven by feedforward neural nets. These models are discriminative: they discriminate categories among sets of images without an explicit model of the image formation process. A more principled way to implement vision (or any data analysis) is to formulate a model of the process that generated the image (the data) and then to invert the process, so as to infer the parameters of the model from the data (for a textbook on this approach in computer vision, see Prince 2012).

For vision, the generative model is an image formation (or graphics) model that generates images from some high-level representation, e.g. a symbolic description of the visual scene. The first challenge is to define such a model. The second challenge is to perform inference on it, i.e. to find the high-level representation that best explains the image, e.g. the maximum a posteriori estimate or the full posterior probability distribution over all possible high-level representations given the image. The idea of an active search for the interpretation that best explains the evidence given our prior knowledge about the world is captured in Helmholtz’s (1866) description of vision as unconscious inference.

Computer vision and biological vision research has always spanned the entire gamut from discriminative to generative approaches (Knill et al. 1996; Yuille & Kersten 2006; Prince 2012). However, the generative approach is practically challenging for computer vision and theoretically challenging for neuroscience. Most computer vision systems, whether using hand-engineered features or deep learning, therefore still rely primarily on discriminative models, learning mostly feedforward computations that process images to produce the desired outputs (but see Prince 2012). In neuroscience, similarly, feedforward models like HMAX (Riesenhuber & Poggio 1999) have been influential.

Image formation involves nonlinear processes such as occlusion. Whereas the inversion of a linear generative model has a closed-form solution that can be implemented in a feedforward computation, inverting a graphics model is computationally much more challenging. Inferring a high-level scene description from an image requires consideration (at some level of abstraction) of a combinatorial explosion of possible configurations of objects and lights.

The deep mystery of vision is exactly how discriminative and generative models are integrated into a seamless and efficient process of inference. Vision might rely on a discriminative feedforward model for rapid recognition at a glance and on recurrent dynamics for iterative refinement of the inference, for correcting the errors of an initial feedforward estimate, or for choosing among a set of hypotheses highlighted by the feedforward pass.

Recurrent neural networks can implement dynamic inference processes of this type and, given recent successes in the domain of language processing, seem poised for a fundamental advance in vision research.

Recurrent computations might converge on a point estimate of the parameters of a generative model of the image. Alternatively, they might implement *probabilistic inference* on a generative model, converging on a representation of the posterior distribution over the generative model’s parameters.

Recurrent message passing can implement *belief propagation*, an algorithm for probabilistic inference on a generative model. If the model captured the causal process giving rise to images, the recurrent dynamics could infer the specific causes (e.g. the objects, their properties, and the lighting) of a particular image. This process can be implemented in recurrent neural networks and might explain how the brain performs optimal cue combination, temporal integration, and explaining away (Lochmann & Deneve 2011).

Belief propagation is a deterministic algorithm for probabilistic inference. Another deterministic proposal is based on probabilistic population codes (Ma et al. 2006). Alternatively, a neural net might perform probabilistic inference by Markov Chain Monte Carlo (MCMC) sampling, using neural stochasticity as a random generator (Hoyer & Hyvarinen 2003; Fiser et al. 2010; Buesing et al. 2011; McClelland 2013; Häfner et al. 2014). A snapshot of neural population activity, in this view, represents a point estimate of the stimulus and a temporal sequence of such snapshots represents the posterior distribution. For near-instantaneous readout of a probabilistic representation several MCMC chains could operate in parallel (Savin & Deneve 2014). The sampling approach naturally handles the representation of joint probability distributions of multiple variables.

These proposals are exciting because they explain how the brain might perform formal probabilistic inference with neurons, linking the biological hardware to the high-level goal of rational information processing. The ultimate goal, of course, is not rational inference, but successful behaviour (survival and reproduction). We should expect the brain to perform probabilistic inference only to the extent that it is expedient to do so in the larger context of successful behaviour.

It remains to be seen how the probabilistic inference proposals of computational neuroscience scale up to the real-world challenges of vision. If they do, they might play a central role in future brain theory and computer vision. The brain clearly handles uncertainty well in many contexts (Tenenbaum et al. 2006; Pouget et al. 2013), so it is helpful to view its inferences as approximations, however rough, to rational probabilistic inference.

At a larger time scale, vision involves top-down effects related to expectation and attentional scrutiny, and active exploration of a scene through a sequence of eye movements, and through motor manipulations of the world. With the recurrent loop expanded to pass through the environment, these processes bring limited resources (the fovea, conscious attention) to different parts of the environment sequentially, selectively sampling the most relevant information while accumulating evidence toward an overall interpretation. Active perception is also being explored in the computational literature. For example, Tang et al. (2014b) built a model for face recognition that uses a convolutional feedforward pass for initialisation and an attentional mechanism for selection of a region of interest, on which probabilistic inference is performed on a generative model, which itself is learned from data.

The challenge ahead is, first, to scale recurrent neural net models for vision to real-world tasks and human performance levels and, second, to fit and compare their representational dynamics to biological brains. Recurrent models are already successful in several domains of AI, including video-to-text description (Venugopalan et al. 2014), speech-to-text recognition (Sak et al. 2014), text-to-text language translation (Sutskever et al. 2014; Cho et al. 2014), and text-to-speech synthesis (Fan et al. 2014). In brain science, recurrent neural network models will ultimately be needed to explain every major function of brain information processing, including vision, other perceptual processes, cognition, and motor control.

## CONCLUSIONS

Computational neuroscience has been very successful by asking what the brain *should* compute (Körding 2007). The normative goals proposed have often led to important insights before being replaced by larger goals. Should the brain efficiently encode sensory information (Barlow 1961)? Or should it infer an accurate probabilistic representation of the world (Barlow 2001)? The ultimate goal is successful behaviour.

Normative theory has driven advances at the cognitive and neuronal levels. Successes of this approach include theories of efficient coding (Barlow 1961, Olshausen & Field 1997, Simoncelli & Olshausen 2001), probabilistic neuronal coding and inference (Hoyer & Hyvärinen 2003, Fiser et al. 2010, Buesing et al. 2011, McClelland 2013, Pouget et al. 2013), Bayesian sensorimotor control (Körding & Wolpert 2006), and probabilistic cognition (Tenenbaum et al. 2006). For low-level sensory representations and for low-dimensional decision and control processes, normative theories prescribe beautiful and computationally simple inference procedures, which we know how to implement in computers and which might plausibly be implemented in biological brains. However, visual recognition and many other feats of brain information processing require inference using massive amounts of world knowledge. Not only are we missing a normative theory that would specify the optimal solution, but, until recently, we were not even able to implement any functioning solution.

Until recently, computers could not do visual object recognition, and image-computable models that could predict higher-level representations of novel natural images did not exist. Deep neural networks put both the task of object recognition and the prediction of high-level neural responses within our computational reach. This advance opens up a new computational framework for modelling high-level vision and other brain functions.

Deep neural net models are optimized for task performance. In this sense, the framework addresses the issue of what the brain *should* compute at the most comprehensive level: that of successful behaviour. In its current instantiation, the deep net framework gives up an explicit probabilistic account of inference, in exchange for neurally plausible models that have sufficient capacity to solve real-world tasks. We will see in the future whether explicitly probabilistic neural net models can solve the real-world tasks and explain biological brains even better.

### Replacing one black box by another?

One criticism of using complex neural networks to model brain information processing is that it replaces one impenetrably complex network with another. We might be able to capture the computations, but we are capturing them in a large net, the complexity of which defies conceptual understanding. There are two answers to the criticism of impenetrability.

First, it is true that our job is not done when we have a model that is predictive of neural responses and behaviour. We must still strive to understand---at a higher level of description---how exactly the network transforms representations across the multiple stages of a deep hierarchy (and across time when the network is recurrent). However, once we have captured the complex biological computations in an artificial neural network, we can study its function efficiently in silico--- with full knowledge of its internal dynamics. Synthetic neurophysiology, the analysis and visualization of artificial network responses to large natural and artificial stimulus sets, might help reveal the internal workings of these networks (Zeiler & Fergus 2014, Girshick et al. 2014, Simonyan et al. 2014, Tsai & Cox 2015, Zhou et al. 2015, Yosinski et al. 2015).

The second answer to the criticism of the impenetrability of neural network models is that we should be prepared to deal with mechanisms that elude a concise mathematical description and an intuitive understanding. After all, intelligence requires large amounts of domain-specific knowledge, and compressing this knowledge into a concise description or mathematical formula might not be possible. In other words, our models should be as simple as possible, but no simpler.

Similar to computational neuroscience, AI began with simple and general algorithms. These algorithms did not scale up to real-world applications, however. Real intelligence turned out to require incorporating large amounts of knowledge. This insight eventually led to the rise of machine learning. Computational neuroscience must follow in the footsteps of AI and acknowledge that most of what the brain does requires ample domain-specific knowledge learned through experience.

### Are deep neural net models similar to biological brains?

The answer to this question is in the eye of the beholder. We can focus on the many abstractions from biological reality and on design decisions driven by engineering considerations and conclude that they are very different. Alternatively, we can focus on the original biological inspiration and on the fact that biological neurons can perform the operations of model units, and conclude that they are similar.

Abstraction from biological detail is desirable and is in fact a feature of all models of computational neuroscience. A model is not meant to be identical to its object, but rather to explain it at an abstract level of description. Merely pointing out a difference to biological brains, therefore, does not constitute a legitimate challenge. For example, the fact that real neurons spike does not pose a challenge to a rate-coding model. It just means that biological brains can be described at a finer level of detail that the model does not address. If spiking were a computational requirement (e.g., Buesing et al. 2011) and a spiking model outperformed the best rate-coding model at its own game of predicting spike rates, or at predicting behaviour, however, then this model would present a challenge to the rate-coding approach.

Many features of the particular type of deep convolutional feedforward network currently dominating computer vision deserve to be challenged in the context of modelling biological vision (see Sidebar *Adversarial examples can reveal idiosyncrasies of neural networks*). The features that deserve to be challenged first are the higher-level computational mechanisms, such as the lack of bypass connections in the feedforward architecture, the lack of feedback and local recurrent connections, the linear--nonlinear nature of the units, the rectified linear activation function, and the max-pooling operation. To challenge one of these features, we must demonstrate that measured neuronal responses or behavioural performance can be more accurately predicted using a model that does not have the feature.

The neural network literature is complex and spans the gamut from theoretical neuroscience to computer science. This literature includes feedforward and recurrent, discriminative and generative, deterministic and stochastic, nonspiking and spiking models. It provides the building blocks for tomorrow’s more comprehensive theories of information processing in the brain. Now that these models are beginning to scale up to real-world tasks and human performance levels in engineering, we can begin to use this modelling framework in brain science to tackle the complex processes of perception, cognition, and motor control.

### The way ahead

We will use modern neural network technology with the goal of approximating the internal dynamics and computational function of large portions of biological brains, such as their visual systems. An important goal is to build models with layers that correspond one-to-one to visual areas, and with receptive fields, nonlinear response properties, and representational geometries that match those of the corresponding primate visual areas. The requirement that the system perform a meaningful task such as object recognition provides a major functional constraint.

Task training of neural networks with millions of labelled images currently provides much stronger constraints than neurophysiological data do on the space of candidate models. Indeed, the recent successes at predicting brain representations of novel natural images are largely driven by task training (Yamins et al. 2014, Khaligh-Razavi & Kriegeskorte 2014, Cadieu et al. 2014). However, advances in massively parallel brain-activity measurement promise to provide stronger brain-based constraints on the model space in the future. Rather than minimizing a purely task-based loss function, as commonly done in engineering, modelling biological brains will ultimately require novel learning algorithms that drive connectivity patterns, internal representations, and task performance into alignment with brain and behavioural measurements.

AI, machine learning, and the cognitive and brain sciences have deep common roots. At the cognitive level, these fields have recently converged through Bayesian models of inference and learning (Tenenbaum et al. 2006). Similar to deep networks, Bayesian nonparametric techniques (Ghahramani 2013) can incorporate large amounts of world knowledge. These models have the advantage of explicitly probabilistic inference and learning. Explaining how such inference processes might be implementated in biological neural networks is one of the major challenges ahead.

Neural networks have a long history in AI, in cognitive science, in machine learning, and in computational neuroscience. They provide a common modelling framework to link these fields. The current vindication in engineering of early intuitions about the power of brain-like deep parallel computation reinvigorates the convergence of these disciplines. If we can build models that perform complex feats of intelligence (AI) and explain neuronal dynamics (computational neuroscience) and behaviour (cognitive science), then--for the tasks tackled -- we will understand how the brain works.

## SUMMARY POINTS

1. Neural networks are brain-inspired computational models that now dominate computer vision and other artificial intelligence applications.
2. Neural networks are networks of interconnected units computing nonlinear functions of their input. Units typically compute linear combinations of their inputs followed by a static nonlinearity.
3. Feedforward neural networks are universal function approximators:
4. Recurrent neural networks are universal approximators of dynamical systems.
5. Deep neural networks stack multiple layers of nonlinear transformations and can concisely represent complex functions like those needed for vision.
6. Convolutional neural nets constrain the input connections of units in early layers to local receptive fields, with weight templates replicated across spatial positions, reducing the number of parameters that need to be learned.
7. Deep convolutional feedforward networks for object recognition are not biologically detailed and rely on nonlinearities and learning algorithms that may fundamentally differ from those of biological brains. Nevertheless they learn internal representations that are highly similar to representations in human and nonhuman primate inferior temporal cortex.
8. Neural networks now scale to real-world artificial intelligence tasks, providing an exciting technological framework for building more biologically faithful models of complex feats of brain information processing.

## FUTURE ISSUES

1. Building systems that engage complex real-world tasks and simultaneously model biological brain activity patterns and behavioural performance (including the overall level of performance, errors, and reaction times or detailed motor trajectories).
2. Increasing biological fidelity in terms of architectural parameters, nonlinear representational transformations, and learning algorithms.
3. Building networks whose layers match the areas of the visual hierarchy in their representational geometries.
4. Predicting behavioural responses to particular stimuli, including similarity judgments and reaction times in discrimination tasks, from neural network representations.
5. Developing supervised learning techniques that drive neural networks into alignment with measured brain activity and behavioural data.
6. Building recurrent neural network models whose representational dynamics resemble those of biological brains.
7. Building neural network models, in which feedforward and recurrent computations interact to implement probabilistic inference on generative models of image formation.
8. Tackling more complex visual functions including categorisation and identification of unique entities, attentional shifts and eye movements that actively explore the scene, visual search, image segmentation, more complex semantic interpretations, and sensory-motor integration.

## TERMS

Unit: model abstraction of a neuron, typically computing a weighted sum of incoming signals, followed by a static nonlinear transformation.
Feedforward network: network whose connections form a directed acyclic graph, precluding recurrent information flow.
Recurrent network: network with recurrent information flow, which produces dynamics and lends itself naturally to the perception and generation of spatiotemporal patterns.
Convolutional network: network where a layer’s preactivation (before the nonlinearity) implements convolutions of the previous layer with a number of weight templates.
Deep neural network: network with more than one hidden layer between input and output layers; more loosely, network with many hidden layers.
Deep learning: machine learning of complex representations in a deep neural network, typically using stochastic gradient descent by error backpropagation.
Universal function approximator: model family that can approximate any function mapping input patterns to output patterns (with arbitrary precision when allowed enough parameters)
Universal approximator of dynamical systems: a model family generating dynamics that can approximate any dynamical system (with arbitrary precision when allowed enough parameters)
Maxpooling: summary operation implementing invariances by retaining only the maxima of sets of detectors differing in irrelevant properties (e.g. local position).
Normalisation: operation (e.g. division) applied to a set of activations so as to hold fixed a summary statistic (e.g. the sum).
Dropout: regularisation method for neural network training with each unit omitted from the architecture with probability 0.5 on each training trial.
Graphics processing unit (GPU): specialised computer hardware developed for graphics computations that greatly accelerates matrix-matrix multiplications and is essential for efficient deep learning.
Supervised learning: learning process requiring input patterns along with additional information about the desired representation or the outputs (e.g. category labels).
Unsupervised learning: learning process that requires only a set of input patterns and captures aspects of the probability distribution of the inputs.
Backpropagation (*and* backpropagation through time): supervised neural-net learning algorithm that backpropagates error derivatives with respect to the weights through the connectivity to iteratively minimise errors.
Generative model: model of the process that generated the data (e.g. the image), to be inverted in data analysis (e.g. visual recognition).
Discriminative model: model extracting information of interest from the data (e.g. the image) without explicitly representing the process that generated the data
Receptive field modelling: predictive modelling of the response to arbitrary sensory inputs of neurons (or measured channels of brain activity).
Representational similarity analysis: method for testing computational models of brain information processing through statistical comparisons of representational distance matrices that characterise population-code representations.
Synthetic neurophysiology: computational analysis of responses and dynamics of artificial neural nets aimed to gain a higher-level understanding of their computational mechanisms.

## ACKNOWLEDGEMENTS

The author thanks Seyed Khaligh-Razavi and Daniel Yamins for helpful discussions, and Patrick McClure and Katherine Storrs for comments on a draft of the manuscript. This research was funded by the UK Medical Research Council (Programme MC-A060-5PR20), by a European Research Council Starting Grant (ERC-2010-StG 261352), and by a Wellcome Trust Project Grant (WT091540MA).

## RELATED RESOURCES

Hinton G (2012) Coursera course “Neural Networks for Machine Learning” https://www.coursera.org/course/neuralnets

Bengio Y, Goodfellow I, Courville A (in progress) “Deep Learning” online book http://www.iro.umontreal.ca/~bengioy/dlbook/

Ng A Coursera course “Machine Learning” https://www.coursera.org/course/ml

Nielson M (in progress) “Neural Networks and Deep Learning” online book http://neuralnetworksanddeeplearning.com/

